# The physical boundaries of public goods cooperation between surface-attached bacterial cells

**DOI:** 10.1101/119032

**Authors:** Michael Weigert, Rolf Kümmerli

## Abstract

Bacteria secrete a variety of compounds important for nutrient scavenging, competition mediation and infection establishment. While there is a general consensus that secreted compounds can be shared and therefore have social consequences for the bacterial collective, we know little about the physical limits of such bacterial social interactions. Here, we address this issue by studying the sharing of iron-scavenging siderophores between surface-attached microcolonies of the bacterium *Pseudomonas aeruginosa*. Using single-cell fluorescent microscopy, we show that siderophores, secreted by producers, quickly reach non-producers within a range of 100 μm, and significantly boost their fitness. Producers in turn respond to variation in sharing efficiency by adjusting their pyoverdine investment levels. These social effects wane with larger cell-to-cell distances and on hard surfaces. Thus, our findings reveal the boundaries of compound sharing, and show that sharing is particularly relevant between nearby yet physically separated bacteria on soft surfaces, matching realistic natural conditions such as those encountered in soft tissue infections.

## 1. Introduction

The study of cooperative interactions in bacteria is of interdisciplinary interest, as it is relevant for understanding microbial community assembly (1,2), the establishment of infections (3–5), and biotechnological processes (6). Bacteria exhibit a wide range of cooperative traits, including the formation of biofilms and fruiting bodies, the secretion of toxins to infect hosts, coordinated swarming, and the scavenging of nutrients from the environment through the secretion of shareable compounds, such as enzymes and siderophores (7,8). While the existing body of work has greatly changed our perception of bacteria – from simple autarkic individuals to sophisticated organisms, interacting and cooperating with each other – there are still considerable knowledge gaps. For instance, many of the insights gained on the sharing of public goods are based on experiments in planktonic batch cultures, where behavioural responses are averaged across millions of cells. This contrasts with the natural lifestyle of bacteria, where individual cells adhere to surfaces and form biofilms, and primarily interact with their immediate neighbours at the micrometre scale (9,10). The mismatch between laboratory and natural conditions has led to controversies in the field regarding the general relevance of microbial cooperation (11–13).

In our paper, we tackle these issues by testing whether and to what extent secreted siderophores are shared between surface-attached individuals of the bacterium *Pseudomonas aeruginosa* using fluorescent microscopy. Siderophores are secondary metabolites produced by bacteria to scavenge iron from the environment, where it typically occurs in its insoluble ferric form or is actively withhold by the host in the context of infections (14,15). In our experiments, we examined the production and secretion of pyoverdine, the main siderophore of *P. aeruginosa* (16), which has become a model trait to study cooperation in bacteria, because it fulfils all the criteria of a cooperative trait: it is costly to produce and secreted outside the cell, where it generates benefits in iron-limited media for the producer itself, but also for nearby individuals with a compatible receptor (17–19). Although highly influential, many of the insights gained are based on batch culture experiments, which tell us little about whether pyoverdine is also shared in surface-attached communities, where molecule diffusion might be limited, and thus the range of sharing constrained (13,20). However, such knowledge is key to understand whether public goods cooperation occurs in natural settings and in infections, where bacteria typically live in biofilms attached to organic and inorganic substrates (8,21).

Here, we present data from fluorescence time-laps microscopy experiments that examined bacterial interactions in real time at the micrometer scale. First, we tested whether pyoverdine molecules, secreted by producing cells, reach individuals that cannot produce pyoverdine themselves but have the receptor for uptake. Such evidence would be a direct demonstration of molecule sharing. Second, we test whether pyoverdine serves as a signalling molecule (22), allowing producers to respond to changes in their social neighbourhood. Specifically, we predict that lower pyoverdine investment is required in a cooperative neighbourhood due to the efficient reciprocal pyoverdine sharing, whereas non-producers, which act as a sink for pyoverdine, should trigger increased investment levels to compensate for pyoverdine loss (23,24). Third, we examined whether pyoverdine diffusivity limits the range across which pyoverdine can be efficiently shared. To this end, we manipulated both the media viscosity, which directly affects molecule diffusion, and the distance between producer and non-producer cells, which increases the diffusion time and reduces the amount of pyoverdine reaching non-producers. Finally, we used time-laps microscopy to quantify fitness effects of pyoverdine production and sharing in growing micro-colonies. Taken together, our experiments shed light on the physical boundaries and individual fitness consequences of public goods sharing.

## 2. Materials and methods

### (a) Strains and media

Our experiments featured the clinical isolate *P. aeruginosa* PAO1 (ATCC 15692), and its clean pyoverdine knock-out mutant (PAO1*ΔpvdD*), directly derived from this wildtype. To be able to distinguish the two strains, we used fluorescent variants of these strains constructed via chromosomal insertion (*att*Tn7::ptac-*gfp, att*Tn7::ptac-*mcherry*) – i.e. PAO1-*gfp*, PAO1-*mcherry*, PAO1Δ*pvdD*-*gfp* and PAO1Δ*pvdD*-*mcherry.* A preliminary experiment revealed that these fluorescent markers did not affect the growth performance of the strains (figure S3). For our gene expression experiments, we used the reporter strain PAO1*pvdA-gfp* (chromosomal insertion: *att*B::*pvdA*-*gfp)* (25). PvdA catalyses an important step in the biosynthesis pathway of pyoverdine (26), and its expression level is therefore a good proxy for the investment into pyoverdine production.

Overnight cultures were grown in 8 ml Lysogeny Broth (LB) medium in 50 ml Falcon tubes, and incubated at 37°C, 200 rpm for ca. 17 hours. Cells were then harvested by centrifugation (3000 rpm/ 3 minutes) and resuspension in 8 ml of 0.8% (saline solution). Subsequently, we diluted the washed cultures in saline solution to an OD = 1 (optical density at 600 nm). For all microscopy experiments, we used CAA medium (per liter: 5 g casamino acids, 1.18 g K_2_HPO_4_*3H_2_O, 0.25 g MgSO_4_*7H_2_O). To create severe iron limitation, we added 450 μM of the iron chelator 2,2-Bipyridin. To create iron-replete conditions, we added 200 μM FeCl_3_. All chemicals were purchased from Sigma-Aldrich (Buchs SG, Switzerland).

### (b) Preparation of microscopy slides

We adapted a method previously described in (27). Standard microscopy slides (76 mm × 26 mm) were washed with EtOH and dried in a laminar flow. We used “Gene Frames” (Thermo Fisher Scientific) to prepare agarose pads. Each frame features a single chamber of 0.25 mm thickness (1.5 x 1.6 cm) and 65 μl volume. The frame is coated with adhesives on both sides so that it sticks both to the microscopy slide and to the cover glass. The sealed chamber is airproof, which is necessary to prevent evaporation and pad deformation during the experiment.

To prepare pads, we heated 20 mL CAA supplemented with agarose (1% unless indicated otherwise) in a microwave. The melted agarose-media mix was subsequently cooled to approximately 50°C. Next, we added the supplements: either 2,2-Bipyridin (450 μM) or FeCl_3_ (200 μM) to create iron-limited or iron-replete conditions, respectively. We pipetted 360 μL of the agarose solution into the gene frame and immediately covered it with a cover glass. The cover glass was pressed down with a gentle pressure to dispose superfluous media. After the solidification of the pad (ca. 30 minutes), we removed the cover glass (by carefully sliding it sideways) and divided the original pad into 4 smaller pads of equal size by using a sterile scalpel. We introduced channels between pads, which served as oxygen reservoir. We then put 1 μL of diluted bacterial culture (OD = 1 cultured diluted by 2.5*10^-4^) on each pad. Two pads were inoculated with a 1:1 mix of pyoverdine producers and non-producers, whereas the other pads were inoculated with a monoculture (either producer or non-producer). After the drop had evaporated, we sealed the pads with a new cover glass. With this protocol, we managed to create agarose pads with consistent properties across experiments.

### (c) Microscopy setup and imaging

All experiments were carried out at the Center for Microscope and Image Analysis of the University Zürich (ZMB) using a widefield Leica DMI6000 microscope. The microscope featured a plan APO PH3 objective (NA = 1.3), an automated stage and auto-focus. For fluorescent imaging, we used a Leica L5 filter cube for GFP (Emission: 480±40 nm, Excitation: 527±30 nm, DM = 505) and a Leica TX2 filter cube for mCherry (Emission: 560±40 nm, Excitation: 645±75 nm, DM = 595). Auto-fluorescence of pyoverdine was captured with a Leica CFP filter cube (Emission: 436±20 nm, Excitation: 480±40 nm, DM = 455). We used a Leica DFC 350 FX camera (resolution: 1392×1040 pixels) for image recording (16 bit colour depth).

### (d) Image processing and blank subtraction

To extract information (cell size, fluorescence) from single cells, images had first to be segmented (i.e. dividing the image into objects and background). Since this is currently a bottleneck for high throughput image analysis (28), we developed a new rapid, reliable and fully automated image segmentation workflow (see supplementary material). The workflow starts with the machine learning, supervised object classification and segmentation tool ilastik (29). Ilastik features a self-learning algorithm that autonomously explores the parameter space for object recognition. We used a low number of phase contrast images from our experiments to train ilastik. Each training round is followed by user inputs regarding segmentation errors. These inputs are then incorporated in the next training round, until segmentation is optimized and error-free. Once the training is completed, batches of microscopy images can be fed to ilastik and segmentation is then executed automatically.

Segmented images were transferred to Fiji, a free scientific image processing software package (30). We wrote specific macro-scripts in Fiji to fully automate the simultaneous analysis of multiple single-cell features such as cell size, shape, fluorescence (see supplementary material for a step-by-step protocol). Next, we applied a pixel-based blank correction procedure in Fiji, to obtain unbiased fluorescence intensities for each cell. For each agarose pad and time point, we imaged four empty random positions on the agarose pad without bacterial cells and averaged the grey values for each pixel. The averaged grey value of each pixel was then subtracted from the corresponding pixel position in images containing cells. This pixel-based blank correction accounts for intensity differences across the field of view caused by the optical properties of the microscope (vignetting). In the experiments where we simultaneously measured *pvdA*-*gfp* expression and pyoverdine fluorescence, we had to further correct for the leakage of pyoverdine signal into the GFP-channel. To do so, we imaged cells of the unmarked wildtype strain, which produced pyoverdine but had no GFP reporter. We measured the pyoverdine signal in the GFP-channel at three different time points (one, three and five hours post-incubation), and then used these values to blank correct the fluorescence intensities in cells with the *pvdA*-GFP reporter.

### (e) Assays measuring *pvdA* expression and pyoverdine fluorescence

To monitor pyoverdine investment by producer cells and pyoverdine uptake by non-producer cells, we quantified natural pyoverdine fluorescence in bacterial micro-colonies in mixed and monocultures over time. For producers, we further measured *pvdA* expression levels over time. Because the excitation wavelength for pyoverdine fluorescence overlaps with the UV range, the high exposure time required to measure natural pyoverdine fluorescence induces phototoxicity. Accordingly, each bacterial micro-colony could only be measured once. To obtain time course data for pyoverdine expression and uptake levels, we thus prepared multiple microscopy slides, as described above, and incubated them at 37 °C in a static incubator. At each time step (one, three and five hours post incubation), we processed two slides for imaging. Exposure time for measuring gfp-fluorescence was 800 ms and for pyoverdine 1500 ms, with a (halogen) lamp intensity of 100%. To guarantee reliable automated image analysis, we only considered positions that were free from non-bacterial objects (e.g. dust) and where all cells laid within one focus layer. We recorded at least five positions per treatment, time point and slide. The experiment was carried out twice, in two independent batches.

### (g) Fitness assays

We used time-laps microscopy to measure the growth performance of pyoverdine producer cells (mCherry-tagged) and non-producer cells (GFP-tagged) in mixed and monoculture. We cut the agarose pad in four patches and inoculated two patches with a 1:1 mix of producers and non-producers, and one patch each with a monoculture. We then chose 20 positions (five per patch) that contained two separated cells (one cell of each strain for mixed cultures and two cells of the same type for monocultures), and imaged these positions sequentially every 15 minutes over 5 hours, using the automated stage function of the microscope. Following a position change, we used the auto-focus function of the microscope in order to keep cells in focus.

We carried out the above fitness assays across a range of different conditions. In a control experiment, we added 200 μM FeCl_3_ to the agarose pad to study strain growth in the absence of iron limitation. Since bacteria grow rapidly in iron-replete media, we stopped the imaging after three hours before micro-colonies started to grow in multiple layers. Next, we monitored strain growth on iron-limited 1% agarose pads supplemented with 450 μM bipyridin. To examine whether pyoverdine sharing and fitness effects depend on the distance between two cells, we performed fitness assays where two cells were positioned: (i) close to one another in the same field of view (average distance between cells 36.21 μM ± 18.17 SD); (ii) further apart in adjacent fields of view (with an estimated minimum distance of 96 μm, given the field of view size of 96 x 128 μm); and (iii) far from one another. This latter condition was created by adding the two strains on opposite ends of an elongated double-sized agarose pad. Finally, we repeated the growth assays in media with increased viscosity using 2 % agarose pads.

### (h) Statistical methods

All statistical analyses were performed in R 3.3.0 (31) using linear models (ANOVA or t-tests). Prior to analysis, we used the Shapiro-Wilk test to check whether model residuals were normally distributed. Since each experiment was carried out in multiple independent experimental blocks, we scaled values within each block relative to the mean of the control treatment (i.e. pyoverdine producer monocultures). For all time-laps growth experiments, we considered the position (i.e. the field of view) as the level of replication. For the analysis of single cell fluorescence data, we considered each cell as a replicate.

## 3. Results

### (a) Pyoverdine diffuses from producers to non-producers

We put mono- and mixed cultures of the wildtype strain PAO1 and its isogenic pyoverdine mutant PAO1*ΔpvdD* (tagged with a fitness-neutral mCherry marker) on iron-limited agarose pads on a sealed microscopy slide. Cultures were highly diluted such that single cells were physically separated from each other at the beginning of the experiment. We then monitored the pyoverdine fluorescence in growing micro-colonies over time for both strains under the microscope. Pyoverdine fluorescence becomes visible in the periplasma, where molecule maturation occurs (13,32) (figure 1b). We found that fluorescence in non-producer colonies was indistinguishable from background signal one hour after incubation, indicating that no detectable pyoverdine had yet been taken up (figures 1a+c and S1). However, pyoverdine fluorescence in non-producer cells significantly increased over time in mixed cultures (LM: F_5,7567_ = 913, p < 0.001) and was significantly higher than the background fluorescence in non-producers growing as monocultures (t-test: t_3945_ = 79.33, p < 0.001, figures 1a+d and S1). This demonstrates that significant amounts of pyoverdine diffuse from producer to non-producer microcolonies even when there is no direct cell-to-cell contact.

**Figure 1.**
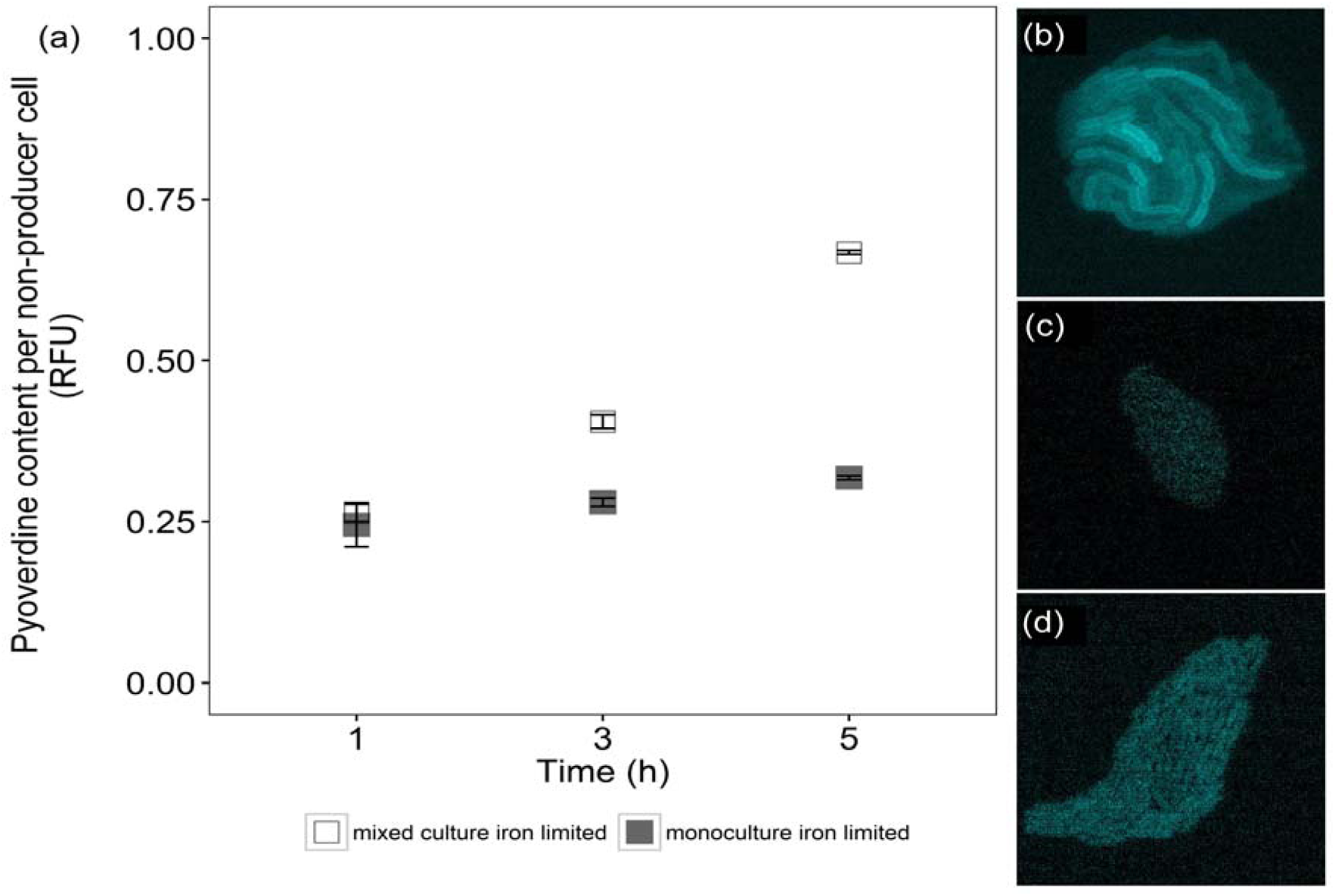
Pyoverdine is taken up by non-producing cells in a time-dependent manner, demonstrating pyoverdine sharing between physically separated, surface-attached micro-colonies. **(a)** Time-course measures on natural pyoverdine fluorescence units (RFU) shows constant background fluorescence in non-producer cells grown in monocultures (filled squares), whereas pyoverdine fluorescence significantly increased in non-producer cells grown in mixed cultures with producers (open squares). Mean relative fluorescence values ± standard errors are scaled relative to producer monocultures after one hour of growth. Representative microscopy pictures show pyoverdine fluorescence in a producer microcolony **(b)**, a non-producer colony from a monoculture **(c)**, and a non-producer colony from a mixed culture **(d)**. Important to note is that only apo-pyoverdine (i.e. iron-free) is fluorescent, and therefore the measured fluorescence intensities represent a conservative measure of the actual pyoverdine content per cell. Furthermore, the fluorescence intensity in producer cells is always higher than in non-producer cells because it represents the sum of pyoverdine uptake and newly synthesized pyoverdine, whereas for non-producers, fluorescence represents pyoverdine uptake only.

### (b) Producers alter pyoverdine investment in the presence of non-producers

To test whether producers respond to changes in their social environment, we followed the expression pattern of *pvdA* (a gene involved in pyoverdine synthesis) and natural pyoverdine fluorescence in growing producer microcolonies (figures 2 and S2). In our control treatment with added iron, both *pvdA* and pyoverdine signal were downregulated compared to iron-limited conditions, demonstrating the functioning and high sensitivity of our reporters. Under iron limitation, meanwhile, *pvdA*-expression was significantly higher in mixed compared to monoculture at one hour (t-test: t_115_ = 5.23, p < 0.001) and three hours (t_860_ = 13.92, p < 0.001) post-incubation (figures 2a and S2a). Pyoverdine fluorescence mirrored *pvdA* expression patterns, with higher pyoverdine levels being detected in producer cells growing in mixed cultures (figures 2b and S2b), although the difference was only significant after three hours (t-test: t_992_ = 13.30, p < 0.001), but not after one hour (t-test: t_88_ = 1.26, p = 0.211). The picture changed five hours post-incubation, where both *pvdA*-expression and pyoverdine fluorescence were significantly lower in mixed compared to monocultures (*pvdA*-expression: t_6441_ = -16.67, p < 0.001; pyoverdine fluorescence: t_6017_ = -50.01, p < 0.001). These analyses demonstrate that producers rapidly alter pyovedine investment in response to the presence of non-producers.

**Figure 2.**
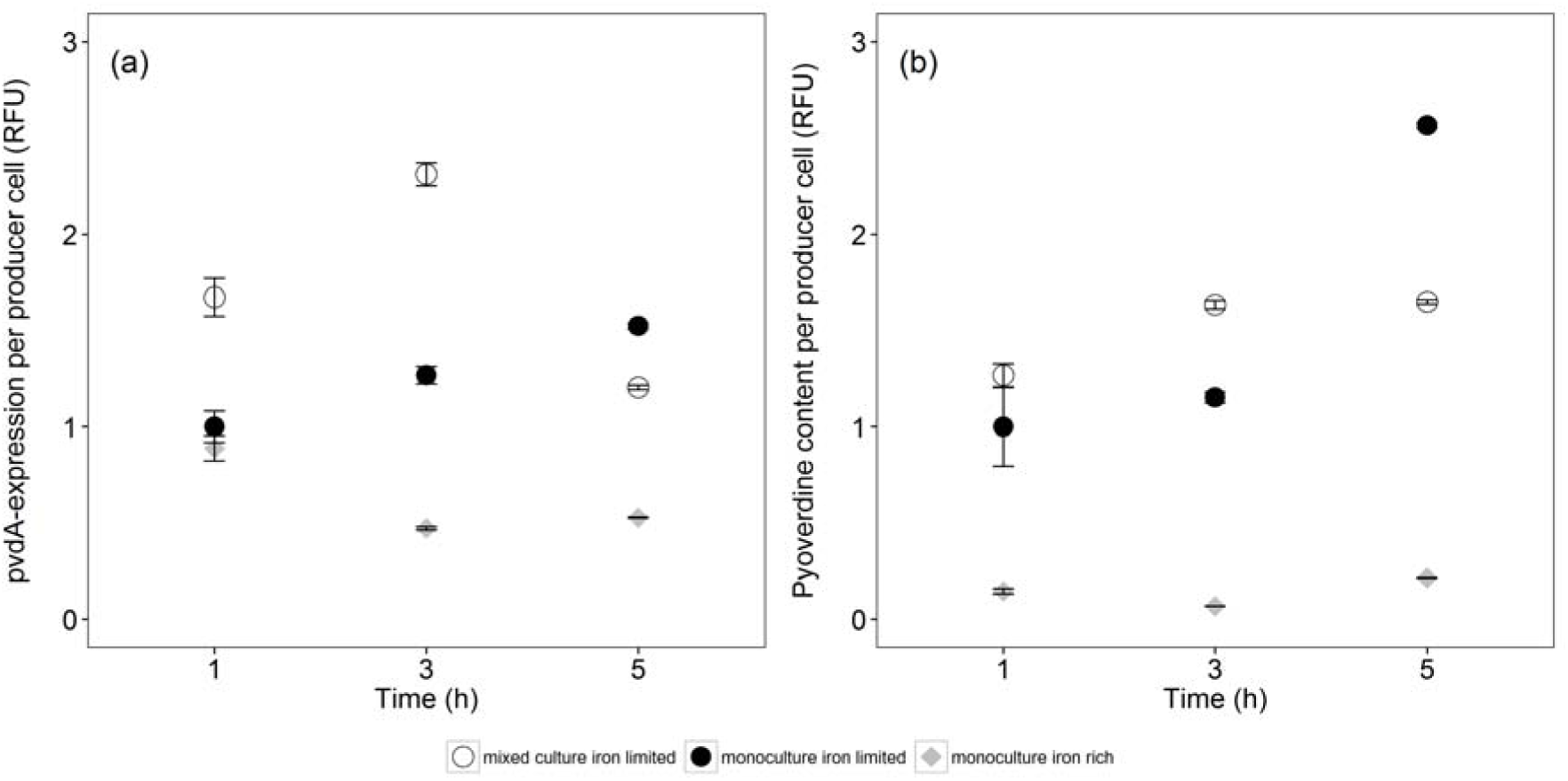
Producer cells adjust their pyoverdine investment level in response to changes in the social environment. **(a)** Time-course data show that *pvdA*, a gene encoding an enzyme involved in pyoverdin synthesis, is down-regulated in iron-rich media (grey diamonds), but up-regulated in iron-deplete media. Importantly, producers exhibited different *pvdA* expression patterns depending on whether they grew together with non-producers (open circles) or as monoculture (filled circles). While producers showed increased gene expression in mixed compared to monoculture after one and three hours, the pattern flipped after five hours. **(b)** The same qualitative pattern was observed when measuring pyoverdine content per cell, as relative fluorescence units (RFU). Fluorescence values are scaled relative to the producer monocultures after one hour of growth. Error bars indicate standard errors of the mean.

### (c) Pyoverdine non-producers outgrow producers in mixed cultures

After having established that pyoverdine is shared between neighbouring, yet physically separated surface-attached microcolonies, we explored the fitness consequences of pyoverdine sharing. This is important because experiments in liquid batch cultures repeatedly revealed that non-producers can outcompete producers, by saving the cost of pyoverdine production, yet exploiting the siderophores produced by others, a phenomenon that is called “cheating” (17,33–36). To examine whether cheating is also possible when bacteria grow as surface-attached microcolonies, we grew producers and non-producers in mono and mixed culture and followed microcolony growth dynamics over time (figure 3). Control experiments in iron-supplemented media revealed that all strains grew equally well regardless of whether they grew in mono or mixed cultures (figure S4). In iron-limited media, however, we found that microcolony growth was significantly reduced for non-producers compared to producers (growth rate: t_23_= -10.57, p < 0.001, figure 3e; cell number: t_23_= -10.27, p < 0.001, figure 3g). This shows that the inability to produce pyoverdine is a major handicap in iron-limited media.

**Figure 3.**
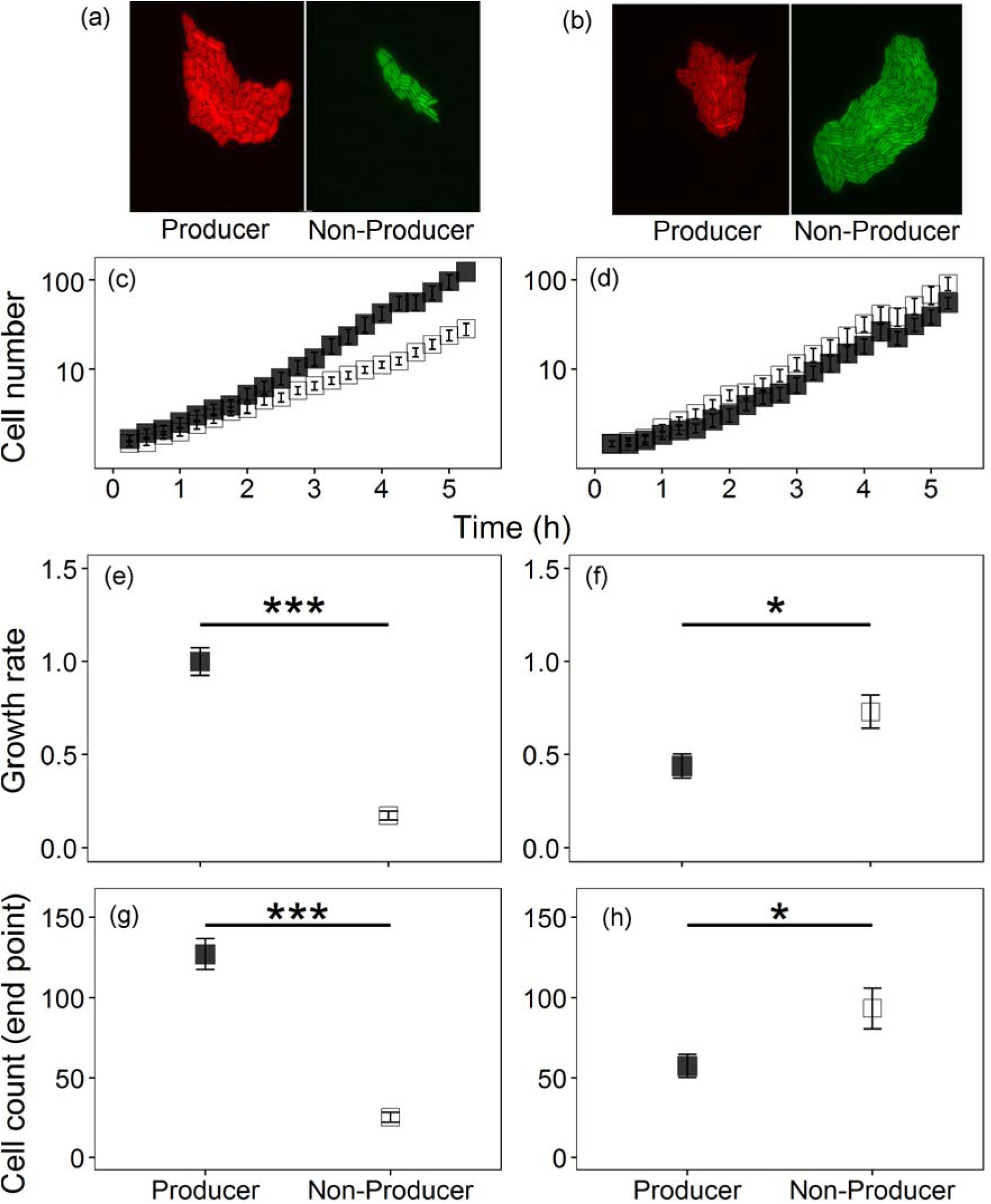
Growth performance of surface-attached microcolonies of pyoverdine producers (filled squares) and non-producers (open squares) in monocultures (left column) and mixed cultures (right column). While pyoverdine non-producers show growth deficiencies in monoculture, due to their inability to scavenge iron, they outcompete the producers in mixed cultures. This growth pattern shows that non-producers save costs by not making any pyoverdine, yet gain fitness benefits by capitalizing on the pyoverdine secreted by the producers. **(a)** and **(b)** show representative microscopy pictures for monocultures and mixed cultures, respectively. The overall growth trajectories of producers and non-producers differ substantially between monocultures **(c)** and mixed cultures (**d)**. While producers had a significantly higher growth rate **(e)** and grew to higher cell numbers **(g)** in monocultures, the exact opposite was the case in mixed cultures for both the growth rate **(f)** and cell number **(h)**. Growth parameters are given relative to the producers in monoculture. Asterisks indicate significant differences and error bars denote standard errors of the mean.

This fitness pattern diametrically flipped in mixed cultures, where non-producer microcolonies grew significantly faster (t_35_= 2.64, p = 0.012, figure 3f) and to higher cell numbers (t_31_ = 2.48, p = 0.019, figure 3h) than producer microcolonies. Intriguingly, non-producers experienced a relative fitness advantage between hours one and three (t-test: t_20_ = 4.53, p < 0.001), but not at later time points (t_22_ = 4.46, p < 0.001; figure S5). This specific period, at which the relative fitness advantage manifests, perfectly matches the timeframe during which producers exhibited highest *pvdA* expression levels, and non-producers started accumulating pyoverdine (figure 2 and S2). Our findings thus provide a direct temporal link between the high costs of pyoverdine investment to producers, the increased benefits accruing to non-producers, and the resulting opportunity for non-producers to act as cheaters and to successfully outcompete producers.

### (d) The physical boundaries of pyoverdine sharing and benefits for non-producers

The above experiments revealed that pyoverdine can be shared between two physically separated microcolonies when grown in the same field of view (128 × 96 μm) under the microscope (average ± SD distance between cells *d* = 36.2 ± 18.2 μm). Next, we asked what the physical limit of pyoverdine sharing is. We thus repeated to above experiment, but this time we focussed on non-producer cells that had no producer cell within the same field of view, but only a more distant producer in an adjacent field of view (minimal distance *d* ~ 100 μm). Under these conditions, we found that non-producers benefited from the presence of more distant producers in the same way as they benefited from the presence of a close producer (figure 4a+b; significantly increased growth of non-producers in mixed culture, for *d* ~ 100 μm, t-test: t_14_ = 4.02, p = 0.001). However, contrary to the previous observation (figure 4a), the producer no longer experienced a significant growth reduction in the presence of a more distant non-producer (figure 4b, for *d* ~ 100 μm, t_9_ = -0.80, p = 0.442). We then expanded the distance between non-producers and producers even further by adding the two strains on opposite ends of a double-sized agarose pad. In contrast to the previous results, these assays revealed that non-producers had significantly lower number of doublings in both mixed (t_13_= -2.41, p = 0.032) and monocultures (t_9_= -4.66, p = 0.001) (figure 4c), showing that pyoverdine diffusion and sharing is disabled across this large distance in the timeframe analysed.

**Figure 4.**
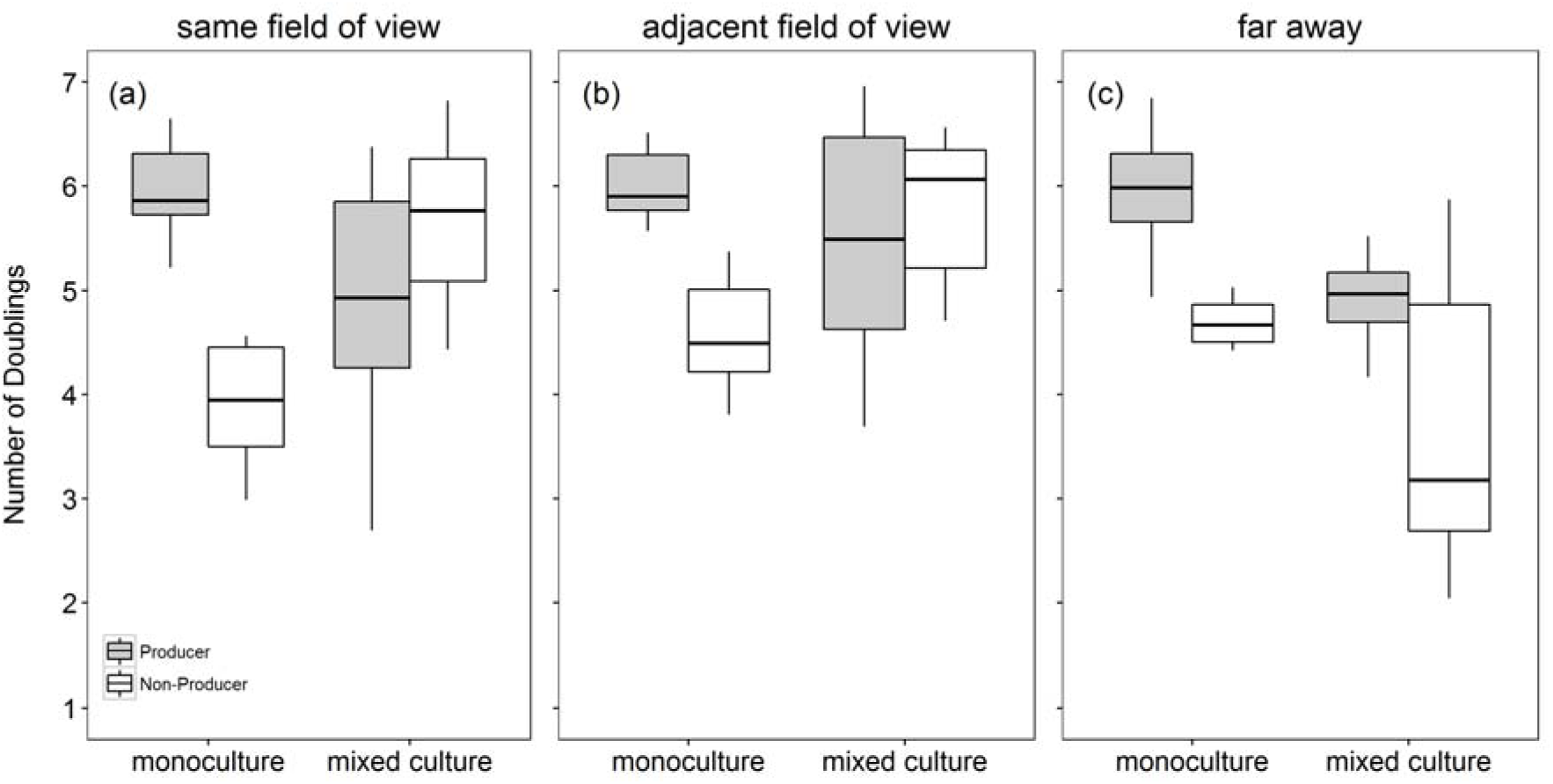
The relative fitness advantage of pyoverdine non-producers in mixed cultures is dependent on the distance between producer (grey) and non-producer (white) microcolonies. In monoculture assays, the non-producers had significantly lower number of doublings than the producers in all experiments. In mixed cultures, meanwhile, the number of doublings of non-producers significantly increased when the producer microcolony was (**a**) within the same field of view (average distance between cells 36 μm), **(b)** in an adjacent field of view (minimal distance ~ 100 μm), but not when producers were far away (on opposite ends of the agarose pad) (**c**). These analyses show that pyoverdine can be shared and exploited across a relative large distance. Boxplots represent the median with 25^th^ and 75^th^ percentiles and whiskers show the 1.5 interquartile range (IQR).

In addition, our microscopy experiment revealed that pyoverdine sharing did not only affect the doubling rate of cells but also their size (figure S6). While non-producer cells were significantly smaller than producer cells in monoculture (LM: F_1,1294_ = 150.90, p < 0.001, measured three hours post-incubation), the cell size of non-producers significantly increased when grown together with a nearby producing neighbour (same field of view *d* ~ 36 μm: t_446_ = 10.24, p < 0.001, figure S6 a; adjacent field of view *d* ~ 100 μm: t_161_ = 4.10, p < 0.001, figure S6 b), but not when producers were far away (on opposite ends of the agarose pad: t_263_= 0.45, p = 0.660, figure S6 c).

While the above experiments examined pyoverdine sharing on 1% agarose pads – a solid yet still moist environment – we were wondering whether pyoverdine sharing is also possible on much harder and drier surfaces. To test this possibility, we repeated the growth experiments on 2% agarose pads. Under these conditions, we observed that non-producers no longer benefited from growing next to producers (no significant difference in the doubling numbers between mono and mixed cultures: t_14_ = -0.98, p = 0.346) (figure 5). This finding is compatible with the view that molecule diffusion is much reduced on very hard surfaces, preventing pyoverdine sharing between adjacent microcolonies.

**Figure 5.**
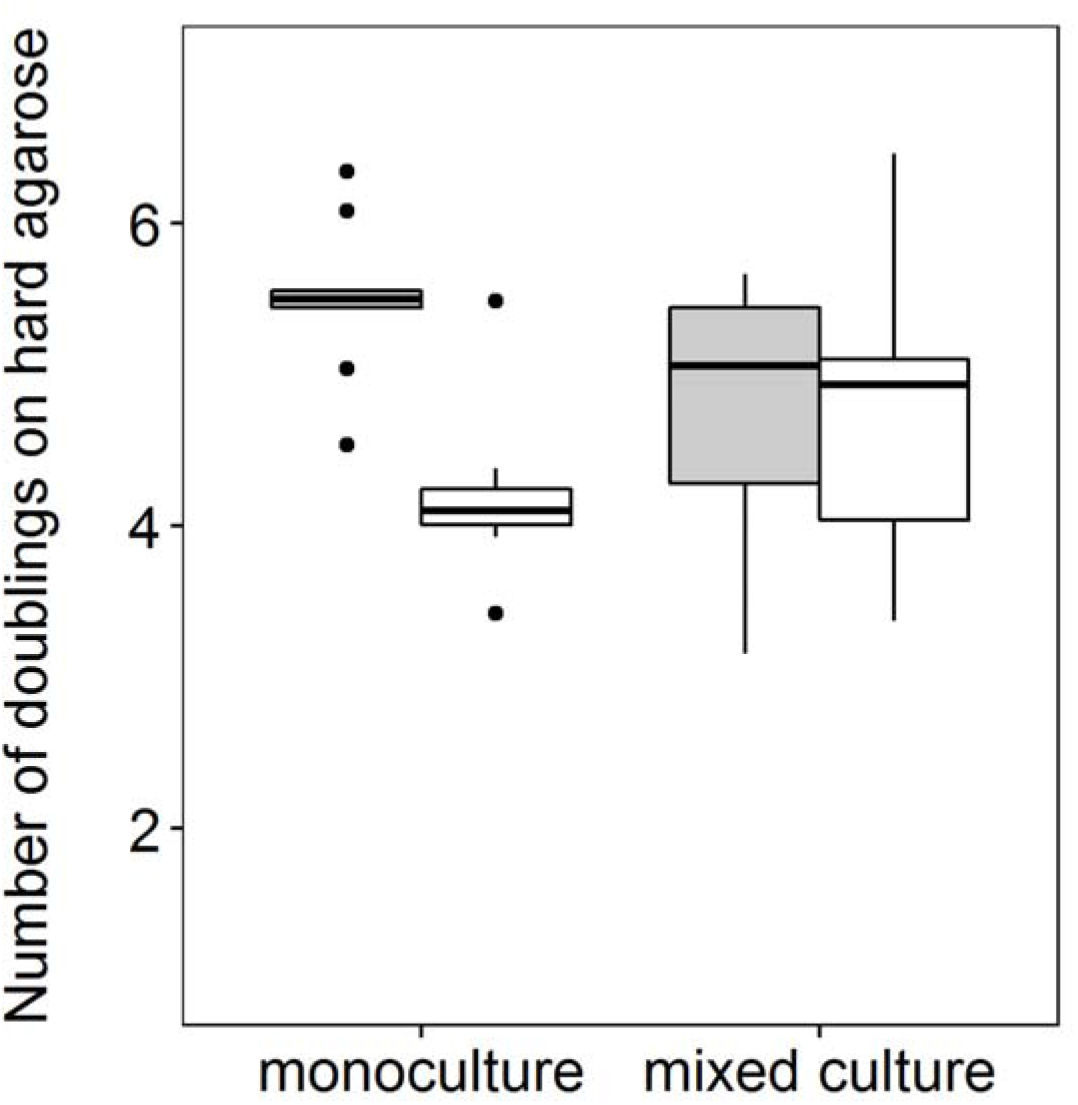
Pyoverdine sharing is impeded on hard surfaces. While the previous experiments showed that pyoverdine is extensively shared between neighbouring microcolonies on relatively soft surfaces (1 % agarose), efficient sharing was no longer possible on hard surfaces (2 % agarose) even when non-producers (open squares) were located next to producers (filled squares). Boxplots represent the median with 25^th^ and 75^th^ percentiles and whiskers show the 1.5 IQR.

## 4. Discussion

Our single-cell analysis on pyoverdine production in *P. aeruginosa* provides several novel insights on the social interaction dynamics between surface-attached bacteria. First, we found that pyoverdine secreted by producer cells is taken up by physically separated non-producer cells, thereby directly demonstrating pyoverdine sharing. Second, we discovered that producer cells rapidly adjust pyoverdine expression levels when non-producers are nearby, by first up-regulating and then down-regulating pyoverdine investment. Third, we demonstrate that pyoverdine sharing has fitness consequences, as it boosts the growth and cell size of non-producers when growing in the vicinity of producers. Finally, we explored the physical limits of pyoverdine sharing and show that on soft surfaces, pyoverdine can be shared across a considerably large scale (at least 100 μM, i.e. ~ 50 times the length of a bacterium), whereas efficient sharing is impeded with larger distances between cells and on hard surfaces. Altogether, our experiments suggest that public goods sharing and exploitation can take place between surface-attached bacteria across a wide range of naturally relevant conditions, and is mediated by molecule diffusion without the need for direct cell-to-cell contact.

Our results oppose previous work claiming that pyoverdine is predominantly shared between adjacent cells within the same microcolony (13). This claim has provoked a controversy on whether pyoverdine, and secreted compounds in general, can indeed be regarded as public goods (12,37). The difference between our experiments and the ones performed by Julou et al. (13) is that their study solely examined pyoverdine content of cells within the same microcolony. Unlike in our study, there was no direct test of whether pyoverdine diffuses to neighbouring microcolonies and what the fitness consequences of such diffusion would be. While we agree that a considerable amount of pyoverdine is probably shared within the microcolony, we here demonstrate that a significant amount of this molecule also diffuses out of the microcolony, providing significant growth benefits to physically separated neighbouring microcolonies. Thus, our work concisely resolves the debate by showing that secreted hydrophilic compounds, such as pyoverdine (38), can be considered as public goods, even in structured environments, with the amount of sharing and the associated fitness consequences being dependent on the distance between neighbouring microcolonies. Moreover, the distance effect we report here at the single-cell level is in line with density effects described at the community level, where secreted compounds are predominantly shared and become exploitable at higher cell densities (i.e. when cell-to-cell distance is reduced 39–42).

A key advantage of single-cell analyses is that it allows the tracking of bacterial behavioural and growth changes in real time with high precision, immediately after the start of an experiment. This contrasts with batch culture experiments, where responses can only be measured after several hours, once the proxies for responses (e.g. optical density) become detectable at the population level. For instance, results from previous batch-culture studies suggest that pyoverdine producers seem to overinvest in pyoverdine when grown together with non-producers (23,24). However, the interpretation of these results based upon a number of assumptions, and the batch-culture approach precluded an in-depth analysis of the temporal pattern and consequences of such overinvestment. Our analysis now provides a nuanced view on the interactions between producers and non-producers. We could show that soon after the inoculation of bacteria on the agarose pad, producers started overexpressing pyoverdine (figures 2 and S2), which coincided with pyoverdine accumulation in non-producer cells (figures 1 and S1), and significant fitness advantages to non-producers (figure S5). Moreover, these findings indicate the that producers can possibly respond to exploitation by down-regulating pyoverdine production at later time points, a response that correlated with the abolishment of further fitness advantages to non-producers.

Our considerations above raise questions regarding the regulatory mechanisms involved in controlling the observed expression changes. Molecular studies suggest that pyoverdine serves as a signalling molecule regulating its own production (22,43). Specifically, when iron-loaded pyoverdine binds to its cognate receptor FpvA, a signalling cascade is triggered, which results in the release of PvdS (the iron-starvation sigma factor, initially bound to the inner cell membrane by the anti-sigma factor FpvR). PvdS then upregulates pyoverdine production. This positive feedback, triggered by successful iron uptake, is opposed by a negative feedback operated by Fur (ferric uptake regulator), which silences pyoverdine synthesis once enough iron has been taken up (16,44). Our results can be interpreted in the light of these feedbacks, given that the relative strength of the opposing feedbacks determines the resulting pyoverdine investment levels (45). For example, producer micro-colonies reach higher cell densities in mono-compared to mixed cultures (figure 3, after 3h: 13.2 ± 2.3 versus 6.7 ± 1.3 cells; after 5h: 122.7 ± 17.9 versus 55.0 ± 8.1 cells, respectively). Higher cell densities likely lead to more efficient pyoverdine sharing, which supposedly stimulates both pyoverdine-signalling and iron uptake. Positive and negative feedback should thus be in balance and result in an intermediate pyoverdine investment levels. Conversely, when producers grow in mixed cultures then cell density is reduced and non-producers serve as a sink for pyoverdine, thereby reducing iron supply to producers. In this scenario, the positive feedback should be stronger than the negative feedback, resulting in the upregulation of pyoverdine. While these elaborations are compatible with the pyoverdine expression patterns observed at hour one and three, the flip in expression patterns between mono and mixed cultures after five hours is more difficult to explain. One option would be that the previously described switch from pyoverdine production to recycling (46–48) occurs earlier in mixed than in monocultures. An alternative option would be that producers can recognize the presence of exploitative cheaters and downscale their cooperative efforts accordingly.

Our results showing that non-producers can outcompete producers in mixed cultures, even when microcolonies are physically separated, confirms predictions from social evolution theory for microbes (49–52). One key condition required for cooperation to be maintained is that cooperative acts must be more often directed towards other cooperators than expected by chance. This interaction probability is measured as the degree of relatedness *r*, a parameter central to inclusive fitness theory (53,54). Traditionally, high relatedness has been associated with the physical separation of cooperators and non-cooperators into distinct patches (54). Our results now show that this traditional view is not necessarily applicable to public goods cooperation in bacteria, because the physical separation of pyoverdine producers and non-producers is insufficient to prevent exploitations and maintain cooperation (figure 3). Clearly, relatedness in our scenario should be measured at the scale at which pyoverdine sharing can occur (50), which exceeds the boundaries of a single microcolony. Thus, in scenarios where microbial cells are immobile, it is the diffusion properties of the public good that determines the degree of relatedness between interacting partners (49,51).

In summary, our finding on pyoverdine sharing and exploitation between physically separated microcolonies has broad implications for our understanding of the social life of bacteria in many natural settings. This is because bacteria typically live in surface-attached communities in aquatic and terrestrial ecosystems, as well as in infections (8,21). Many of these natural habitats feature soft surfaces, as mimicked by our experimental set up, making the diffusion and sharing of secreted compounds between cells highly likely. However, our work also reveals physical limits to public goods cooperation, namely on hard surfaces, where public good diffusion and sharing is impeded. This shows that whether or not a secreted compound is shared is context-dependent (38), and relies, amongst other factors, on the physical properties of the environment.

## Data Archiving Statement

Upon acceptance, raw data and the code for single cell analysis will be made available on Dryad.

## Contributions

MW and RK developed the experimental methods. MW carried out the experiments. MW and RK carried out the statistical analysis. MW and RK drafted the manuscript and all authors gave final approval for publication.

## Acknowledgments

We thank Urs Ziegler and Caroline Aemisegger for help with the microscope, Moritz Kirschmann for advice regarding single cell data analysis.

